# Odorant receptor orthologues in conifer-feeding beetles display conserved responses to ecologically relevant odors

**DOI:** 10.1101/2022.02.22.481428

**Authors:** Rebecca E. Roberts, Twinkle Biswas, Jothi Kumar Yuvaraj, Ewald Grosse-Wilde, Daniel Powell, Bill S. Hansson, Christer Löfstedt, Martin N. Andersson

**Affiliations:** Department of Biology, Lund University, Lund, Sweden; Department of Evolutionary Neuroethology, Max Planck Institute for Chemical Ecology, Jena, Germany; Faculty of Forestry and Wood Sciences, Czech University of Life Sciences Prague, Czech Republic; Global Change Ecology Research Group, School of Science, Technology and Engineering, University of the Sunshine Coast, Sippy Downs, QLD, Australia

**Keywords:** Functional characterization, de-orphanization, HEK293 cells, evolutionary conservation, Coleoptera, Curculionidae

## Abstract

Insects are able to detect a plethora of olfactory cues using a divergent family of odorant receptors (ORs). Despite the divergent nature of this family, related species frequently express several evolutionarily conserved OR orthologues. In the largest order of insects, Coleoptera, it remains unknown whether OR orthologues have conserved or divergent functions in different species. Using HEK293 cells, we addressed this question through functional characterization of two groups of OR orthologues in three species of the Curculionidae (weevil) family, the conifer-feeding bark beetles *Ips typographus* L. (‘Ityp’) and *Dendroctonus ponderosae* Hopkins (‘Dpon’) (Scolytinae), and the pine weevil *Hylobius abietis* L. (‘Habi’; Molytinae). The ORs of *H. abietis* were annotated from antennal transcriptomes. Results show highly conserved response specificities, with one group of orthologues (HabiOR3/DponOR8/ItypOR6) responding exclusively to 2-phenylethanol (2-PE), and the other group (HabiOR4/DponOR9/ItypOR5) responding to angiosperm green leaf volatiles (GLVs). Both groups of orthologues belong to the coleopteran OR subfamily 2B, and share a common ancestor with OR5 in the cerambycid *Megacyllene caryae*, also tuned to 2-PE, suggesting a shared evolutionary history of 2-PE receptors across two beetle superfamilies. The detected compounds are ecologically relevant for conifer-feeding curculionids, and are probably linked to fitness, with GLVs being used to avoid angiosperm non-host plants, and 2-PE being important for intraspecific communication and/or playing a putative role in beetle-microbe symbioses. To our knowledge, this study is the first to reveal evolutionary conservation of OR functions across several beetle species and hence sheds new light on the functional evolution of insect ORs.

## Introduction

Insects are able to detect thousands of different chemical cues that convey important information about the environment, including plant volatiles, microbial odors, and pheromones (Dahanukar et al. 2005; Hansson & Stensmyr 2011; Kandasamy et al. 2019; Stensmyr et al. 2012). This sophisticated discernment of odors is possible via large suites of odorant receptors (ORs), which bind and detect odor molecules in peripheral olfactory sensory neurons (OSNs), triggering neuronal signals to be processed by the central nervous system (Brand et al. 2018; Clyne et al. 1999; Sato et al. 2008; Vosshall et al. 1999; Wicher et al. 2008). These seven-transmembrane proteins form heteromeric complexes with a highly conserved co-receptor (Orco), which is necessary for odor responses in most insects by contributing to the formation of a ligand-gated ion channel (Butterwick et al. 2018; Larsson et al. 2004).

The OR gene family evolves according to a birth-and-death model, in which duplication events represent the birth of genes, and deletion or pseudogenization their death (Nei et al. 2008). Accordingly, OR genes are often found in tandem arrays on insect chromosomes, with significant variation in the size of OR repertoires between species (Andersson et al. 2015; Benton 2015; Brand & Ramírez 2017). Frequently, the majority of ORs in a given species are present within species- or taxon-specific phylogenetic OR-lineage radiations (Mitchell et al. 2020). Within such radiations, novel olfactory functions may evolve due to relaxed constraints or positive selection post gene duplication, provided the duplicated gene is retained and expressed (Andersson et al. 2015; Hou et al. 2021). Despite the general divergent nature of this receptor family, certain ORs are conserved across species, with simple (1:1) orthologous OR-pairs typically being present among related species. However, such clear orthology is usually rare or absent when comparing more distantly related insect species from different families, which has been shown in e.g., beetles (Coleoptera) (Mitchell & Andersson 2020; Mitchell et al. 2020). Whether OR orthologues share the same or similar olfactory functions, or if functions have diverged in different species, has been studied primarily in Lepidoptera and Diptera (e.g., Bohbot et al. 2011; M. Guo et al. 2021). Such studies are important for advancing our understanding of the functional evolution of the insect OR family, as they may inform shared ecological relevance of certain compounds in different species, hence shared selection pressures acting on the olfactory sense of insects. For example, the host- and oviposition cues 1-octen-3-ol, indole and skatole, respectively, are detected by orthologous groups of ORs across several mosquito species (Dekel et al. 2016; Ruel et al. 2019). In moths, some olfactory functions are widely conserved among OR orthologues, including both the detection of specific plant odors and sex pheromone compounds, whereas other orthologues are functionally different (Gonzalez et al. 2015; H. Guo et al. 2021; M. Guo et al. 2021).

In contrast, nothing is known about the functions of OR orthologues in beetles, which is not surprising given the very few ORs that have been functionally characterized in this large order. The response profiles of seven ORs have been characterized from the Eurasian spruce bark beetle *Ips typographus* L. (‘Ityp’; Curculionidae) (Hou et al. 2021; Roberts et al. 2021a; Yuvaraj et al. 2021), two ORs from the red palm weevil *Rhynchophorus ferrugineus* Olivier (Curculionidae) (Antony et al. 2021; Ji et al. 2021), one OR from the dark black chafer *Holotrichia parallela* Motschulsky (Scarabaeidae) (Wang et al. 2020), one OR from the Adonis ladybird *Hippodamia variegata* Goeze (Coccinellidae) (Xie et al. 2022) and three ORs from the hickory borer *Megacyllene caryae* Gahan (‘Mcar’; Cerambycidae) (Mitchell et al. 2012). Among the characterized ORs of this cerambycid, McarOR5 responded strongly to the male-produced pheromone component 2-phenylethanol (2-PE) (Mitchell et al. 2012).

McarOR5 belongs to the beetle OR subfamily named Group 2B, which contains conserved OR lineages with receptors from several beetle families, including the large family of true weevils, Curculionidae (Mitchell et al. 2020). This beetle family harbors the damaging conifer-feeding bark beetles of the Scolytinae subfamily and many other weevils that are pests of agriculture and forestry, such as the pine weevil *Hylobius abietis* L. (‘Habi’; Molytinae) (Shin et al. 2018). The 2-PE is ecologically relevant for several conifer-feeding beetles. For instance, it is part of the attractive odor bouquets released by the fungal symbionts of *I. typographus* (Kandasamy et al. 2019) and the compound has also been identified from the hindgut of male beetles, where it may be produced by yeasts (Leufvén et al. 1984), in highest amounts before the acceptance of females (Birgersson et al. 1984). The compound is the primary odorant for one of the characterized OSN classes of *I. typographus* (Kandasamy et al., 2019), suggesting that this species is likely to have an OR tuned to this compound. Also *Dendroctonus* bark beetles produce this compound (Sullivan 2005), including the mountain pine beetle *D. ponderosae* Hopkins (‘Dpon’; Curculionidae), in which 2-PE reduces the attraction to the aggregation pheromone (Pureswaran et al. 2000). In the pine weevil *H. abietis*, 2-PE operates as a strong anti-feedant present in deterrent non-host plants (Eriksson et al. 2008). Interestingly, 2-PE is also produced by gut bacteria of *H. abietis*, and the behavior of this species, to cover their laid eggs with feces and frass containing deterrent compounds, may protect the eggs from being eaten by conspecifics (Axelsson et al. 2017; Borg-Karlson et al. 2006).

Due to the widespread use of 2-PE in the ecologies of conifer-feeding curculionids, we hypothesized that the compound may be detected by evolutionarily and functionally conserved ORs and that these receptors may be related to McarOR5. To test this hypothesis, we used HEK293 cells to functionally characterize two clades with simple OR orthologues from three curculionids (*I. typographus, D. ponderosae*, and *H. abietis*), i.e., the orthologues in OR Group 2B that are positioned closest to McarOR5 in the OR phylogeny. The OR repertoires of the two bark beetles have been previously reported (Andersson et al. 2013; Andersson et al. 2019; Yuvaraj et al. 2021); however, to obtain the OR sequences from *H. abietis* we sequenced, analyzed, and annotated male and female antennal transcriptomes. Our results show that 2-PE is indeed detected by functionally conserved and highly specific OR orthologues in all three curculionids. These ORs share a common ancestor with McarOR5 suggesting functional conservation also across two beetle superfamilies (Curculionoidea and Chrysomeloidea). Additionally, green leaf volatile (GLV) alcohols, abundant in non-host angiosperm plants and generally avoided by conifer-feeding beetles (Zhang & Schlyter 2004), were detected by the second assayed clade of curculionid ORs. These receptors share evolutionary history with the ORs detecting 2-PE. Altogether, our findings suggest strong functional conservation in ORs detecting ecologically important odors, and thereby expand our knowledge of the functional evolution of the OR family in the largest order of insects.

## Materials and Methods

### *Sequencing, assembly, annotation, and analyses of the* Hylobius abietis *antennal transcriptome*

Wild beetles were collected by hand at Balungstrand’s sawmill in Enviken, close to Falun in mid-Sweden, and kindly provided by Prof. Göran Nordlander (Swedish University of Agricultural Sciences, Uppsala, Sweden). The antennae from 20 males and 20 females were excised and collected separately in tubes kept on dry ice and then stored at −80 °C. The antennae were homogenized using Tissue-tearor model 98370-365 (Bartlesville, OK, USA), and total RNA was isolated using the RNeasy Minikit (Qiagen, Hilden, Germany). Extractions yielded 4.8 μg and 3.1 μg of high-quality total RNA from male and female antennal samples, respectively. These RNA samples were used for both transcriptome sequencing and molecular cloning.

The RNA samples were DNase-treated and then underwent poly-A enrichment and library construction using a TruSeq v2 Library Preparation Kit (Illumina, San Diego, CA, USA), followed by 150 bp paired-end sequencing on an Illumina HiSeq 3000 platform at the Max Planck-Genome center (Cologne, Germany). The sequencing produced 36,455,021 and 35,770,733 paired-end reads from the male and female samples, respectively. Adaptor sequences and low-quality reads were removed using Trimmomatic (v 0.36) (Bolger et al. 2014) with a custom screening database, before performing *de novo* assemblies with Trinity version 2.4.0 (Grabherr et al. 2011). Reads from males and females were assembled separately, and also combined. Contigs from the Trinity output were clustered to reduce the number of redundant transcripts using CD-HIT-EST (v 4.6.8) (Li & Godzik 2006) with a sequence identity threshold of 0.98. Primarily, the sex-combined non-redundant assembly was used for downstream annotation of OR-encoding transcripts, and was comprised of 46,669 predicted protein-coding ‘genes’ with their respective isoforms and together with other non-coding genes totaled 199,035 transcripts. The average transcript length was 824 bp with an N50 of 1,570 bp. The completeness of the sex-combined assembly was firstly assessed using the Benchmarking Universal Single-Copy Orthologs (BUSCO v5.2.2; https://busco.ezlab.org/) tool performed against the Insecta odb10 dataset, including 1,367 reference genes (Manni et al. 2021). This analysis revealed 96.6% complete (C) BUSCOs, of which 59.8% were present as single copy genes (S) and 40.2% as duplicated genes (D). Only 30 (2.3%) BUSCOs were missing (M) from the assembly and 17 (1.3%) BUSCOs were fragmented (F), indicating the majority of transcripts were represented and were full length. Mapping of the clean reads to the non-redundant assembly resulted in an overall alignment rate of 94.06%, further demonstrating a high level of completeness for this assembly. The RNAseq reads have been deposited in the SRA database at NCBI under the BioProject accession number PRJNA783427.

*H. abietis* OR genes were annotated through exhaustive tBLASTn searches against the assemblies at an *e*-value cut-off at 1.0. The OR query sequences were obtained from *I. typographus* (Andersson et al. 2013; Yuvaraj et al. 2021), *D. ponderosae* (Andersson et al. 2013; Andersson et al. 2019), *Anoplophora glabripennis* (McKenna et al. 2016), *M. caryae* (Mitchell et al. 2012), and *Leptinotarsa decemlineata* (Schoville et al. 2018). All annotated HabiOR sequences were included in additional tBLASTn searches against the *H. abietis* assemblies until all novel OR hits were exhausted. Except for the HabiOrco gene which was only assembled to full length in the male-specific assembly, all OR genes were annotated from the sex-combined assembly, and no OR genes were uniquely found or assembled to higher completeness in the two sex-specific assemblies. A few partial OR genes could be extended by joining overlapping transcripts with identical sequences. The names of these genes were given a ‘JOI’ suffix according to established nomenclature (Andersson et al. 2019, and references therein). Likewise, OR genes missing the N-terminus or C-terminus were given the suffixes ‘NTE’ and ‘CTE’, respectively, to their names. Single letter abbreviations were used in combinations (i.e., J, N, C) for genes with multiple suffixes. Transcripts encoding partial OR sequences below 170 amino acids and those that that did not overlap with other OR sequences in multiple sequence alignments were discarded as they may not represent unique genes. Likewise, for ORs sharing >96% amino acid identity, only one transcript was kept in the dataset to exclude potential assembly isoforms or allelic variants. The identified HabiOR genes were given names from HabiOR1 to HabiOR78 following their groupings in the OR phylogeny (Fig. 1), with consecutive numbering within the major coleopteran OR clades (Mitchell et al. 2020).

**Figure 1.**
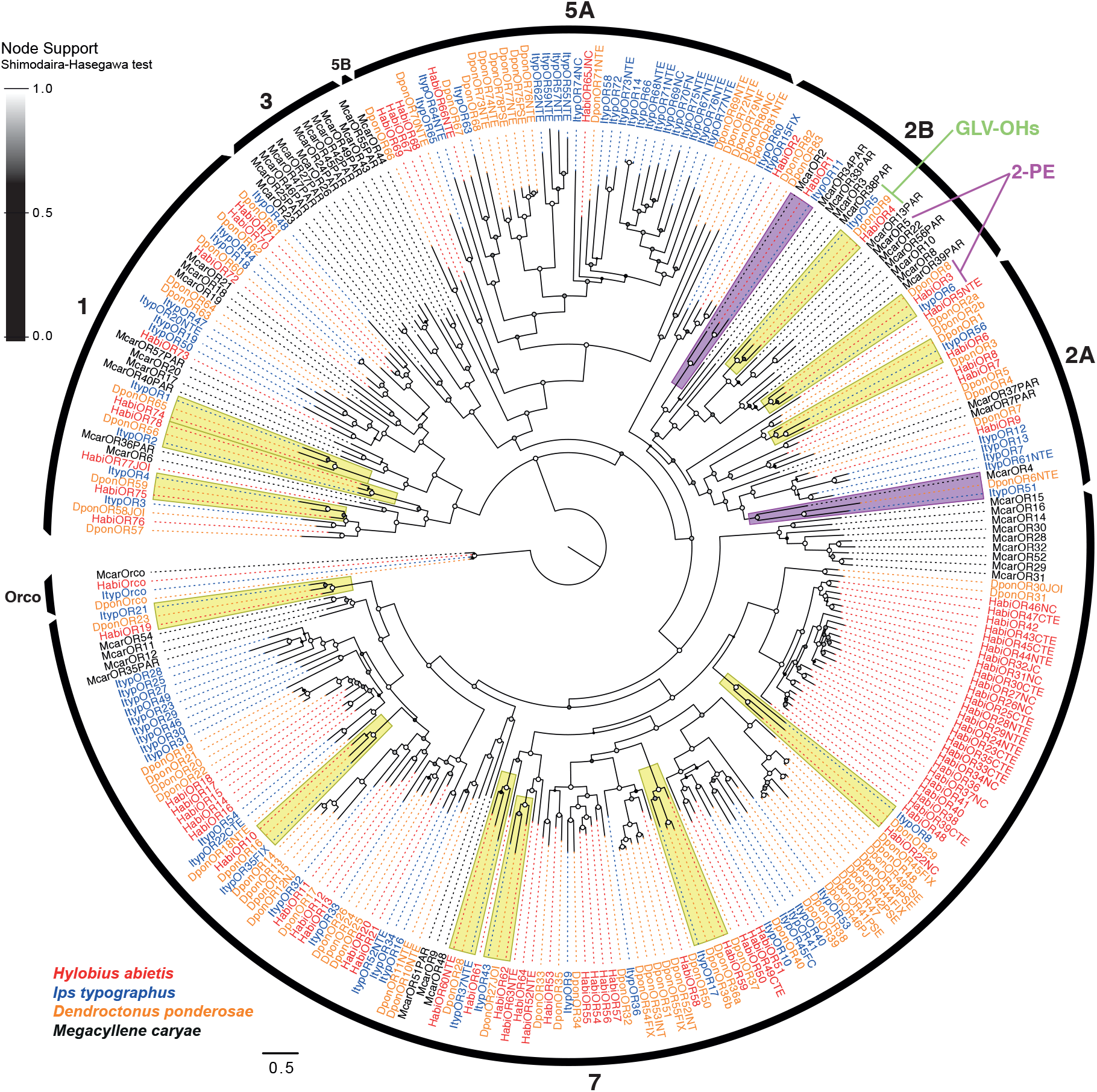
Maximum likelihood phylogeny of odorant receptors (ORs) from beetles of Curculionidae and Cerambycidae. Included are OR amino acid sequences from the curculionids *Hylobius abietis* (‘Habi’; red), *Ips typographus* (‘Ityp’; blue), *Dendroctonus ponderosae* (‘Dpon’; orange), and the cerambycid *Megacyllene caryae* (‘Mcar’; black). The tree is based on a MAFFT alignment, constructed using FastTree, and rooted with the conserved Orco lineage. The major coleopteran OR groups are indicated by the black arcs (Mitchell et al. 2020). The twelve groups of simple (1:1:1) OR orthologues across the three curculionids are highlighted in yellow; two clades housing putative simple orthologues shared by the cerambycid *M. caryae* and two of the curculionids are highlighted in purple. Receptors in OR group 2B that were functionally characterized in the present study and the previously characterized McarOR5 (Mitchell et al. 2012) are labeled by the compounds that activate them (GLV-OHs = green leaf volatile alcohols; 2-PE = 2-phenylethanol). Local node support values were calculated using the Shimodaira-Hasegawa (SH) test implemented in FastTree, and are indicated on branch nodes by the shaded circles; support increases with the brightness of the circles. The sources of sequence data and explanation of receptor suffixes are detailed in the Materials and Methods section.

To analyze the expression levels of OR genes in male and female antennae, clean reads were mapped to the ORFs of annotated HabiOR genes using the align_and_estimate_abundance.pl script from the Trinity v2.4.0 software package (Haas et al. 2013) with default parameters except for --est_method RSEM--aln_method bowtie2 --trinity_mode. The rationale for mapping to the ORFs of OR genes, and not to all transcripts in the assembly, was because some OR transcripts contained misassembled fragments in non-coding regions, which could bias the estimated expression level of the OR gene.

The HabiOR amino acid sequences were aligned with the OR sequences from *D. ponderosae* (Andersson et al. 2019), *I. typographus* (Yuvaraj et al. 2021), and *M. caryae* (Mitchell et al. 2012) using MAFFT v7.450 (Katoh et al. 2002; Katoh & Standley 2013), implemented in Geneious Prime v.2020.0.5 (Biomatters Ltd. Auckland, New Zealand). The alignment of a few partial OR sequences were corrected manually. Three miss-aligned partial McarOR sequences (McarORs 41PAR, 50PAR, and 53PAR) were excluded from analysis since their alignments could not be corrected with confidence. Uninformative regions were excised using trimAl v.1.2 (Capella-Gutiérrez et al. 2009) with the following settings: similarity threshold 0, gap threshold 0.7, and minimum 25% conserved positions. The trimmed alignment was used to construct a phylogenetic tree of ORs using FastTree v.2.1.11 at default settings (Price et al. 2010). Local node support values were calculated using the Shimodaira-Hasegawa (SH) test implemented within FastTree. The tree was rooted with the Orco lineage, and color coded in FigTree v.1.4.3 (Rambaut 2014). Final graphical editing was performed using Adobe Illustrator.

### Molecular cloning and generation of HEK293 cell lines

The protocols for cloning, cell line generation, and culturing have been previously described (Andersson et al. 2016; Corcoran et al. 2014; Yuvaraj et al. 2021). Briefly, OR (and Orco) genes with added 5’ Apa1 and 3’ Not1 restriction sites, *cacc* Kozak sequence, and N-terminal epitope tags (Myc for Orco, V5 for ORs) were ligated into the pcDNATM4/TO (Orco) or pcDNATM5/TO (ORs) mammalian expression vectors (Thermo Fisher Scientific). For OR genes that required codon optimization for functional expression in HEK293 cells (Roberts et al. 2021a), the nucleotide sequences were submitted to the Thermo Fisher Scientific GeneArt Portal and codon optimized for *Homo sapiens*, excluding the start methionine and the epitope tag, then synthesized and ligated into the pcDNATM5/TO expression vector. Because we had no access to biological material from *D. ponderosae*, the two functionally assayed DponOR genes were synthesized as codon optimized versions directly, and also because several wildtype beetle OR genes are not functionally expressed in HEK cells (Roberts et al. 2021a). A codon optimized gene of HabiOR4 was also tested since Western blots (below) failed to detect this protein from cells transfected with the wildtype gene. Receptors encoded by wildtype genes that were detected by Western blot were not codon optimized because they all showed band intensities similar to, or higher than, several ORs previously characterized in this system (Yuvaraj et al. 2018; Yuvaraj et al. 2021). Sequences of codon optimized genes from the two species are provided in Supplementary Table 1. Sequences of OR genes cloned from antennal cDNA (ItypOR5, ItypOR6, HabiOR3, and HabiOR4) have been deposited in GenBank under the accession numbers OL865310-OL865313.

OR genes in expression vectors were transformed into HB101 ampicillin-resistant competent cells (Promega), plated on ampicillin-containing agar, and incubated overnight at 37 °C. Resulting colonies were PCR-screened with vector-specific primers, spread onto a new plate, and incubated at 37 °C for 4-6 hours. Positive colonies were sub-cultured overnight at 37°C in LB broth containing ampicillin. Plasmid DNA from the overnight cultures was harvested via extraction with the PureLinkTM HiPure Plasmid Filter Midiprep kit (Thermo Fisher Scientific), and insert sequence was confirmed by Sanger sequencing at the on-site DNA Sequencing Facility (Dept. Biology, Lund University) using the BigDye^®^ Terminator v1.1 Cycle Sequencing Kit (Thermo Fisher Scientific). Positive plasmids were linearized with FspI, PciI, or BstZ17I (New England Biolabs, Ipswich, MA, United States) restriction enzymes and incubated overnight at 37 °C.

Linearized plasmids containing the ItypOrco gene were transfected into HEK293 cells containing a tetracycline-inducible repressor (TREx) using Lipofectamine 2000 (Thermo Fisher Scientific) (Corcoran et al. 2014). Twenty-four hours post-transfection, antibiotics (blasticidin for TREx, zeocin for Orco; both NEB) were added to select successfully transfected cells. Once a stable cell line was generated, protein expression of the ItypOrco gene was confirmed via Western blot analysis and functionality of Orco was confirmed using the Orco agonist VUAA1 (described below) (Jones et al. 2011). We recently found that the DponOrco gene is not functionally expressed in HEK cells (Roberts et al. 2021a); we therefore co-expressed the DponOR genes with the ItypOrco gene to allow functional characterization of these ORs. For consistency, the HabiOR genes were also tested together with the ItypOrco. Due to the conserved function of Orco and the relatedness among the three beetle species, we assumed that this strategy would not affect the OR response specificities. This assumption is supported by previous studies showing that combinations of OR and Orco proteins from different taxa assemble into functional receptor complexes responding properly to known ligands (Corcoran et al. 2018). The linearized OR gene-containing plasmids were transfected into the stably expressing TREx/ItypOrco cell line as described above, and cultured with the antibiotic hygromycin (Gold BioTech) to select successfully transfected cells. Resulting cell lines were frozen at −80 °C before functional assays.

### Protein extraction and Western blot analysis

Cells were cultured without antibiotics for 24 hours before the expression of Orco and OR genes was induced with doxycycline (Sigma). At 16 hours post-induction, cells were pelleted via centrifugation and total protein extraction was performed as described by Corcoran et al. (2014). Protein extractions from non-induced cells served as negative controls. Western blot was performed using 25 μg of total protein from each sample and standard protocols for mixed molecular weight proteins. Primary antibodies (rabbit anti-Myc for Orco, rabbit anti-V5 for ORs) were added at a ratio of 1:2000, and the secondary antibody (anti-rabbit +IgG, HRP-linked for both Orco and ORs) was added at a ratio of 1:5000 (all antibodies from Cell Signaling Technology), as described by Andersson et al. (2016).

### Functional characterization of odorant receptors

Ligand-binding activity of cell lines co-expressing Orco and ORs was tested via a calcium fluorescence assay using a CLARIOstar Omega plate reader (BMG Labtech, Ortenberg, Germany) according to previously described protocols (Andersson et al. 2016; Corcoran et al. 2014; Yuvaraj et al. 2021). Briefly, cells were plated in black-walled poly-D-lysine coated 96-well plates (Corning Costar) and incubated overnight. Cells in half the wells were treated with doxycycline to induce expression of exogenous Orco and OR genes 16 hours prior to testing, leaving the non-induced cells as negative controls. The calcium-sensitive fluorophore Fluo-4AM (Life Technologies) was loaded into all wells, and plates were incubated in the dark at room temperature for 30 min before being washed with assay buffer, then incubated for another 30 min in the dark at room temperature prior to the assay.

The test odor panel included 62 compounds (Supplementary Table 2) that are ecologically relevant for conifer-feeding curculionids, including pheromone compounds, volatiles from conifers and angiosperms as well as odorants produced by bark beetle fungal symbionts (Kandasamy et al. 2019; Kandasamy et al. 2021; Yuvaraj et al. 2021). The Orco agonist VUAA1 was tested (at 50 μM) on each cell line as a positive control for functional Orco expression. Test odors were diluted in DMSO and assay buffer with a final concentration of 30 μM in the plate wells for screening experiments. Compounds were tested over a minimum of three biological replicates (plates), and were pipetted into three induced and three non-induced wells per plate, creating three technical replicates per plate. A negative control of 0.5% DMSO in assay buffer (vehicle) was tested on each cell line. Background fluorescence was measured for both induced and non-induced cells immediately before compounds were added to the wells, with ligand-binding response of cells measured as the percentage increase in fluorescence from background readings 10 s post stimulation. Mean responses of cells to the added ligands were calculated using GraphPad Prism 6 (GraphPad Software Inc., La Jolla, CA, United States).

A response of ≥1% increase in fluorescence in induced cells was required for a compound to be considered active, provided a significantly higher response in induced compared to non-induced cells. Hence, a general linear model (GLM) with “induction” as a fixed factor and “plate” as a random factor (to account for inter-plate variation) was performed using IBM SPSS statistics v.25. Bonferroni correction to maintain the α-level at 0.05 (for up to 12 multiple comparisons within a cell line) was undertaken to avoid reporting false positives (Type I statistical error). Compounds eliciting an increase in fluorescence of 3% or more at the 30 μM screening concentration were tested in subsequent dose-response experiments. Half-maximal effective concentrations (EC_50_) were calculated using the non-linear curve fit regression function in GraphPad Prism (version 6). Calculations of EC_50_ values were only performed for compounds with (reasonably) sigmoid dose-response curves.

## Results

### HabiOR annotation, expression, and phylogenetic analysis of ORs

The HabiOrco gene and 78 HabiOR genes were annotated from the antennal transcriptome assembly, of which 51 transcripts encoded full length proteins. The 28 partial HabiORs encompassed 174 to 397 amino acids. Three of the ORs were extended by joining overlapping sequences from two different transcripts. Annotation details and sequences of the HabiORs are presented in Supplementary Table 3. Sequence reads from the male and female samples were mapped to the open reading frames (ORFs) of annotated OR genes for estimation of relative OR gene expression levels. This analysis showed that the HabiOrco gene is clearly more highly expressed than any of the 78 HabiORs (Supplementary Table 3). The ORs showed a range of expression levels (from 0.03% to 4.2% of the Orco expression in males; from 0% to 5.7% in females), with no specific OR standing out as being particularly highly expressed compared to the others. Expression was similar in the two sexes; the only putative exceptions being the partial HabiOR27NC and HabiOR46NC that showed twice the expression in females compared to males, and HabiOR48 with twice the expression in males compared to females. Expression levels of the functionally characterized HabiOR3 and HabiOR4 were intermediate, with a somewhat higher estimate for HabiOR4 in both sexes.

Recently, a phylogenetic analysis of the ORs across several coleopteran superfamilies defined and revised nine main monophyletic groups of ORs (Mitchell et al. 2020). Our phylogenetic analysis including ORs from the Curculionidae and Cerambycidae shows that the distribution of HabiORs among the nine OR groups is similar to that of the other two curculionids (*I. typographus* and *D. ponderosae*) in the analysis (Figure 1), with most (55) ORs located within Group 7, followed by Group 1 (9 ORs), Group 5A and 2A (5 ORs in each), and 2B (4 ORs). The main differences between *H. abietis* and the two bark beetle species are the stronger bias of HabiORs towards Group 7, including a large HabiOR-radiation of 26 receptors (HabiOR23-HabiOR48), and comparatively few HabiORs in Group 5A. The latter may be explained by the generally poor antennal expression of Group 5A ORs (Yuvaraj et al. 2021); indeed, the vast majority of Group 5A ORs from *D. ponderosae* was not found in the initial transcriptome analyses (Andersson et al. 2013), but later recovered from the genome (Andersson et al. 2019). As with other curculionids, our analysis indicates that *H. abietis* entirely lacks ORs from Groups 3, 4, 5B, and 6. Similar to our previous study, the OR phylogeny did not recapitulate the monophyly of Group 2B, which is likely explained by the few species and narrow taxonomic range sampled in this study (Yuvaraj et al. 2021).

Our OR phylogeny suggests 12 highly supported clades of simple (1:1:1) OR orthologues conserved across the three curculionids (Figure 1). We also found that two of the McarORs appear to have representative orthologues in at least some curculionids, i.e., McarOR2 grouped together with HabiOR1 and ItypOR11, and McarOR4 with DponOR6NTE and ItypOR51 (Figure 1). Within OR Group 2B, two orthologous groups of curculionid ORs (ItypOR5/DponOR9/HabiOR4 and ItypOR6/DponOR8/HabiOR3) were positioned close to McarOR5, responding to 2-PE -a pheromone component in this species (Figure 1). Hence, these six ORs were targeted for functional characterization to investigate whether evolutionarily related beetle ORs within and between coleopteran superfamilies may have the same response specificities. The amino acid identities between ItypOR5, DponOR9, HabiOR4 range from 50.5% to 57.8%, and for ItypOR6, DponOR8, and HabiOR3 between 60.0% and 67.5%.

### Conserved responses to green leaf volatiles and 2-phenylethanol in OR orthologues

The HEK cells transfected with each of the six above-mentioned beetle OR genes were analyzed for OR protein detection using Western blots. Except for HabiOR4, the OR proteins were clearly detected, and only from cells induced to express the exogenous receptor genes, demonstrating proper regulation by the repressor system (Supplementary Figure 1). Gene sequences codon-optimized for expression in human cells were used for the DponORs because we had no access to biological material from this species (Roberts et al. 2021a). Additionally, because the HabiOR4 protein was not detected from cell lines transfected with the wildtype OR gene, this gene was also codon-optimized and used in functional assays. The superscript HsCO (*Homo sapiens* Codon Optimized) was added to the names of codon optimized OR genes.

In the OR phylogeny, the orthologous curculionid ORs HabiOR4, ItypOR5, and DponOR9 group most closely to the 2-PE receptor in the cerambycid *M. caryae* (McarOR5; Figure 1). Screening experiments testing 62 ecologically relevant compounds at 30 μM concentration showed that neither of these curculionid ORs responded to 2-PE. Instead, HabiOR4^HsCO^ responded to five six-carbon green leaf volatile (GLV) alcohols, abundant in angiosperm non-host plants, with significantly stronger responses in induced *vs*. non-induced cells (Figure 2A). *Z*3-hexenol elicited the strongest response in the screening experiments (7.3% increased fluorescence; F_1,14_ = 183.8; p < 0.001), followed by slightly weaker and similar responses to each of *E*2-hexenol, *Z*2-hexenol, and 1-hexanol (5.2-5.7%; F_1,14_ = 48.9-118.8; all p < 0.001), and a yet weaker response to *E*3-hexenol (3.6%; F_1,14_ = 40.5; p < 0.001). Subsequent dose-response experiments largely recapitulated the screening data in terms of response magnitudes elicited by the five compounds at the higher concentrations (Figure 2B). Nevertheless, these experiments indicated similar sensitivities to the four most active ligands (EC_50_ values: *Z*3-hexenol – 7.99 μM, *E*2-hexenol – 3.88 μM, 1-hexanol – 2.85 μM, and *Z*2-hexenol – 7.34 μM), whereas the sensitivity to *E*3-hexenol was lower as shown by its weaker response at most tested concentrations and the non-sigmoid shape of the dose-response curve (EC_50_ could not be estimated).

**Figure 2.**
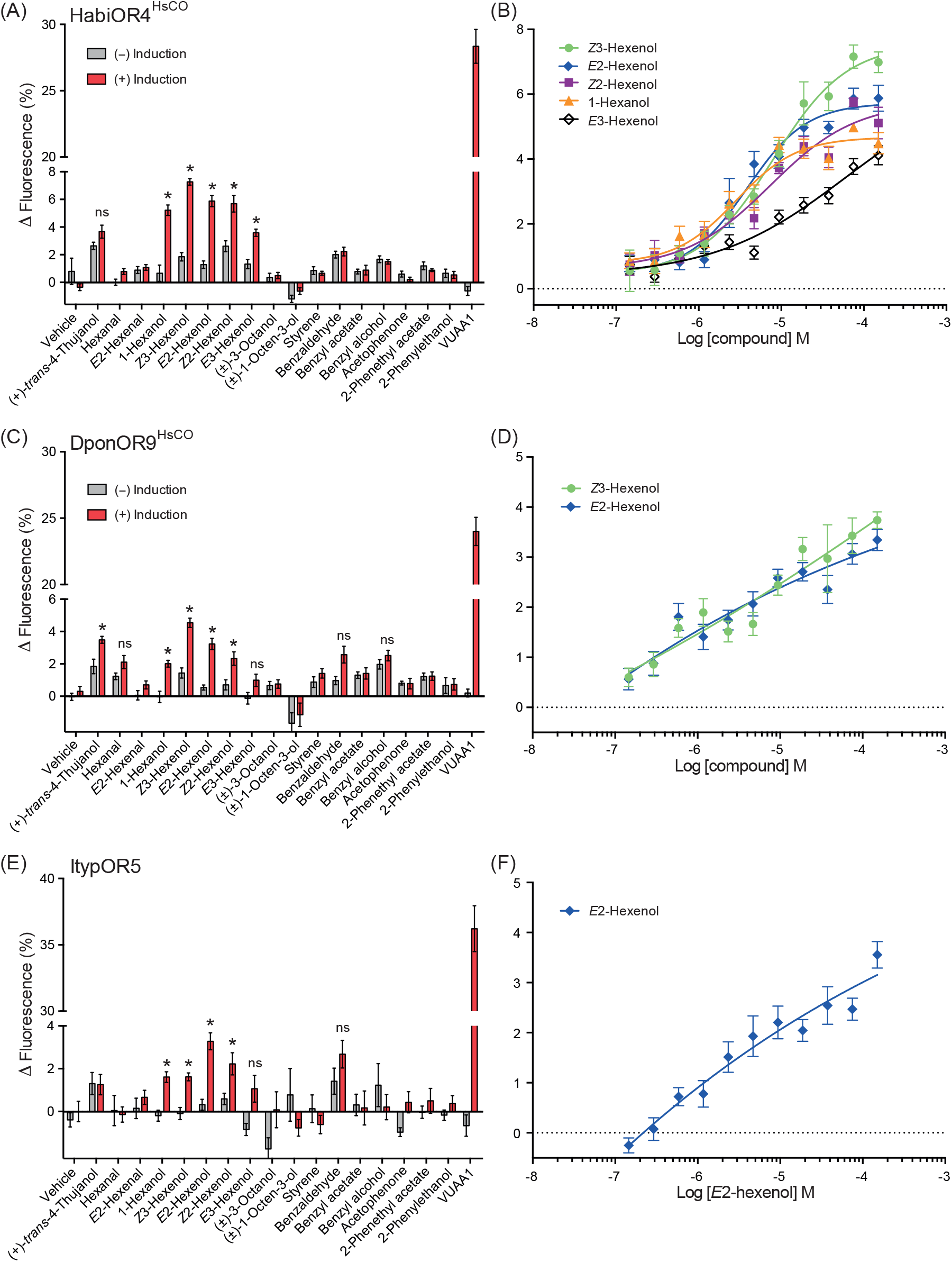
Conserved responses to green leaf volatile (GLV) alcohols in curculionid odorant receptor (OR) orthologues. **(A)** Response of *Hylobius abietis* OR4 (*Homo sapiens* codon optimized; HabiOR4^HsCO^) to select compounds in the screening experiments (30 μM stimulus concentration; *n* = 3 biological replicates, *n_total_* = 9). **(B)** Dose-dependent response of HabiOR4^HsCO^ to the five active GLVs, indicating similar sensitivities to four of the compounds (see main text for EC_50_ values; *n* = 3-5 biological replicates, *n_total_* = 9-15). **(C)** Screening responses of *Dendroctonus ponderosae* OR9 (*H. sapiens* codon optimized; DponOR9^HsCO^; *n* = 3 biological replicates, *n_total_* = 9), and **(D)** dose-dependent responses of DponOR9^HsCO^ to the two most active ligands (*n* = 5 biological replicates, *n_total_* = 15). **(E)** Screening response of *Ips typographus* OR5 (ItypOR5; *n* = 3-5 biological replicates, *n_total_* = 9-15), and **(F)** dose-dependent response of ItypOR5 to the most active ligand (*n* = 6 biological replicates, *n_total_* = 18). Ligand-induced activation was recorded from cells induced (+) to express the exogenous Orco and OR genes and from non-induced (-) control cells. VUAA1 was tested at 50 μM as a control for functional Orco expression. Asterisks indicate significantly stronger responses in induced compared to non-induced cells (at p < 0.001; see main text for details on statistics). Ligands eliciting < 3% increased fluorescence in screening assays were excluded from dose-response trials. Error bars show SEM. All data from induced and non-induced cells to all 62 test compounds are reported in Supplementary Data 1.

A similar response profile was apparent for DponOR9^HsCO^ (Figure 2C). Again, *Z*3-hexenol elicited the highest response in the screening experiments (4.5%; F_1,14_ = 51.5; p < 0.001), followed by *E*2-hexenol (3.2%; F_1,14_ = 44.2; p < 0.001), *Z*2-hexenol (2.3%; F_1,14_ = 48.3; p < 0.001), and 1-hexanol (2.0%; F_1,14_ = 24.6; p < 0.001). The slightly increased fluorescence seen upon stimulation with *E*3-hexenol was not statistically significant after correction for multiple statistical comparisons. In contrast to HabiOR4, (+)-*trans*-4-thujanol elicited a significant response in DponOR9^HsCO^ (3.5%; F_1,14_ = 17.0; p = 0.001); however, half of this response was also evident in the non-induced control cells suggesting that factors unrelated to the DponOR contributed to the cells’ response. The two GLV compounds eliciting responses above 3% increased fluorescence in the screening were further examined in dose-response trials, showing somewhat stronger responses to *Z*3-hexenol as compared to *E*2-hexenol at the higher concentrations but similar responses at intermediate and lower concentrations (Figure 2D). EC_50_ values could not be estimated due to the non-sigmoid shape of the dose-response curves, which is commonly seen for ORs with comparatively low response magnitudes (Roberts et al. 2021a).

ItypOR5 responded to the same four GLV alcohols as did DponOR9^HsCO^, albeit with overall lower response magnitudes (Figure 2E), which is in accordance with this OR being detected as a fainter band on Western blot (Supplementary Figure 1) compared to the other orthologues (see also Roberts et al. 2021a). The rank order between compounds was slightly different, with *E*2-hexenol eliciting the highest response (3.3%; F_1,24_ = 39.8; p < 0.001), followed by *Z*2-hexenol (2.2%; F_1,23_ = 19.3; p < 0.001), and similar responses to *Z*3-hexenol (1.6%; F_1,24_ = 27.9; p < 0.001) and 1-hexanol (1.6%; F_1,24_ = 30.5; p < 0.001). As with DponOR9^HsCO^, the slightly increased fluorescence elicited by *E*3-hexenol in induced cells was not statistically significant after correction for multiple comparisons. *E*2-hexenol activated ItypOR5 in a dose-dependent manner (Figure 2F; EC_50_ could not be estimated due to the shape of the dose-response curve); the other GLVs were not assayed in dose-response experiments due to their screening responses being below 3%.

Compared to the GLV-responding ORs, the orthologous receptors HabiOR3, ItypOR6, and DponOR8 are positioned a bit further away from McarOR5 in the OR phylogeny; yet they are part of the same OR clade (Figure 1). The three ORs all responded exclusively to 2-PE, with significantly stronger responses in induced compared to non-induced cells. The highest response was recorded for HabiOR3 (10.5% increased fluorescence; F_1,19_ = 324.7; p < 0.001; Figure 3A), followed by ItypOR6 (5.7%; F_1,14_ = 85.9; p < 0.001; Figure 3E), and DponOR8^HsCO^ (4.0%; F_1,14_ = 80.4; p < 0.001; Figure 3C). Each of these receptors responded to 2-PE in a dose-dependent manner with estimated EC_50_ values at 8.54 μM, 3.45 μM, and 5.12 μM for HabiOR3, ItypOR6, and DponOR8^HsCO^, respectively (Fig. 3B, D, F).

**Figure 3.**
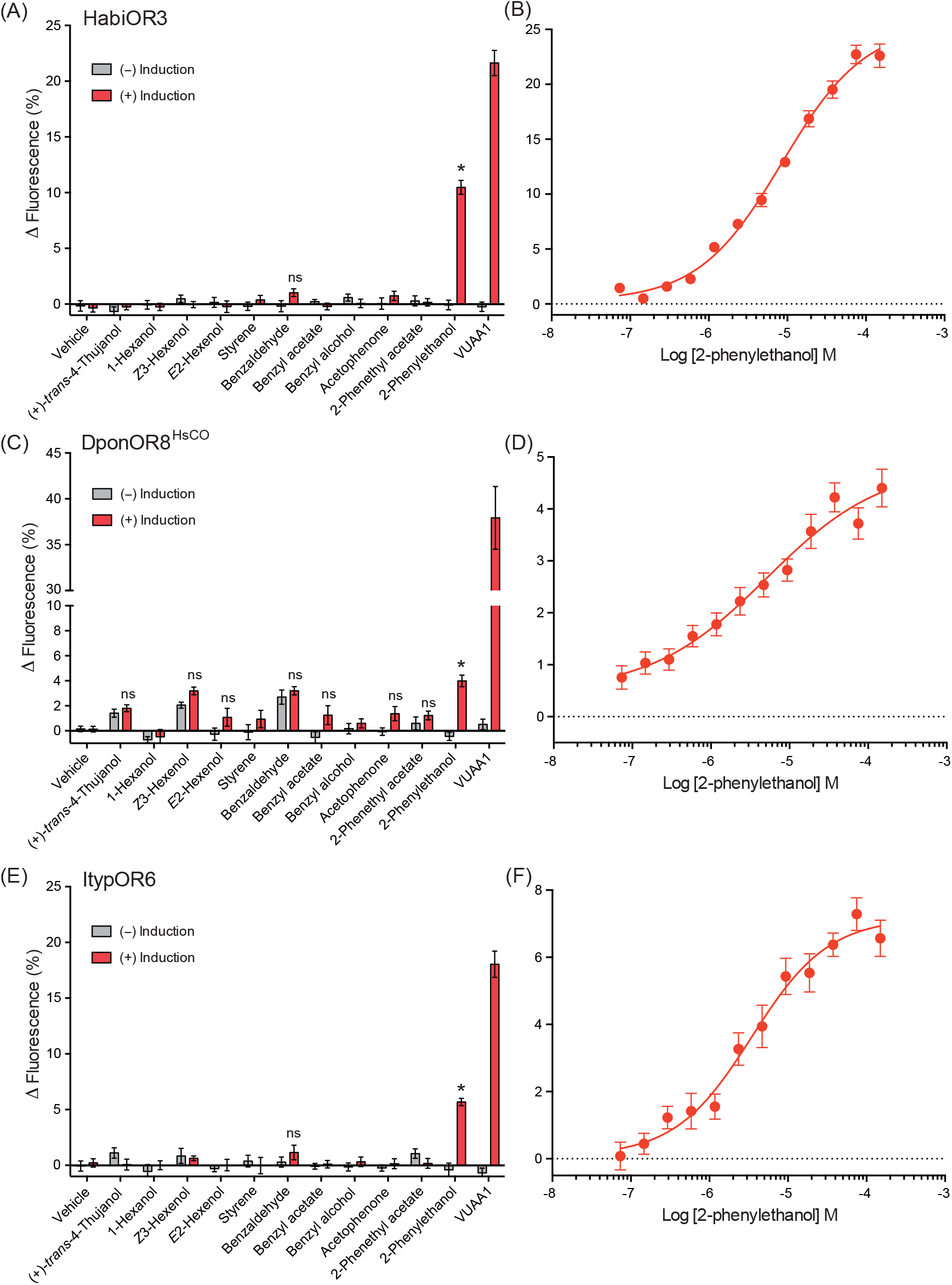
Conserved responses to 2-phenylethanol in curculionid odorant receptor (OR) orthologues. **(A)** Response of *Hylobius abietis* OR3 (HabiOR3) to select compounds in the screening experiments (30 μM stimulus concentration; *n* = 3-4 biological replicates, *n_total_* = 9-12). **(B)** Dose-dependent responses of HabiOR3 (*n* = 3 biological replicates, *n_total_* = 9; see main text for EC_50_ value). **(C)** Screening responses of *Dendroctonus ponderosae* OR8 (*H. sapiens* codon optimized; DponOR8^HsCO^; *n* = 3 biological replicates, *n_total_* = 9), and **(D)** dose-dependent responses of DponOR8^HsCO^ (*n* = 9 biological replicates, *n_total_* = 27; see main text for EC_50_ value). **(E)** Screening responses of *Ips typographus* OR6 (ItypOR6; *n* = 3 biological replicates, *n_total_* = 9), and **(F)** dose-dependent responses of ItypOR6 (*n* = 3 biological replicates, *n_total_* = 9; see main text for EC_50_ value). Ligand-induced activation was recorded from cells induced (+) to express the exogenous beetle Orco and OR genes and from non-induced (-) control cells. VUAA1 was tested at 50 μM as a control for functional Orco expression. Asterisks indicate significantly stronger responses in induced compared to non-induced cells (at p < 0.001; see main text for details on statistics). Error bars show SEM. All data from induced and non-induced cells to all 62 test compounds are reported in Supplementary Data 1.

## Discussion

To allow for functional characterization of *H. abietis* ORs, antennal transcriptomes were sequenced and the ORs annotated, suggesting a similarly sized OR repertoire (79 ORs) as in *I. typographus* (73 ORs) and *D. ponderosae* (86 ORs) (Andersson et al. 2019; Yuvaraj et al. 2021). The largest number of HabiORs was found in OR Group 7, and *H. abietis* stands out by its large OR-radiation within this group. This differs from both the bark beetles and the red palm weevil *R. ferrugineus*, but displays similarity with the large radiation of Group 7 ORs in the sweetpotato weevil *Cylas formicarius* (Antony et al. 2016; Bin et al. 2017). Our previous study indicated that the two scolytine bark beetles *I. typographus* and *D. ponderosae* share 17 highly-supported simple OR orthologues (Yuvaraj et al. 2021). Here, we included also the pine weevil from the Molytinae subfamily in the analysis, and found that 12 simple orthologues are conserved across these three species. The observation of fewer orthologues in this broader analysis is consistent with the general positive correlation between species relatedness and occurrence of OR orthology (Mitchell et al. 2020). Our OR phylogeny also support the notion that OR orthology is rare across beetle superfamilies (Mitchell & Andersson 2020; Mitchell et al. 2020).

To investigate whether OR orthologues are functionally conserved in the three curculionids, and whether evolution of OR functionality may be traced beyond the beetle superfamily level, we characterized two curculionid OR clades within beetle OR subfamily 2B, both of which are evolutionarily related to the cerambycid 2-PE receptor McarOR5 (Mitchell et al. 2012). Our results show conserved functions across the tested OR orthologues, with one group (HabiOR3/DponOR8/ItypOR6) responding exclusively to 2-PE, even though several structurally similar compounds also were tested (Supplementary Table 2). The other group (HabiOR4/DponOR9/ItypOR5) responded to several six-carbon GLV alcohols. Our results further show that the 2-PE receptors in the curculionids share a common ancestor with McarOR5, which is also the case for their GLV-responding ORs. Interestingly, the GLV receptors in the curculionids are the most closely related to McarOR5, while their 2-PE receptors are part of a sister clade. This suggests that the response to 2-PE may be ancestral and that new functions such as GLV-responsiveness have subsequently evolved (neofunctionalization) in the more recent OR lineages of the curculionids. Another possibility would be that the ancestral receptor was broadly tuned to both 2-PE and GLVs and that the more recent OR lineages, derived from gene duplication, may have evolved higher specificity for either GLVs or 2-PE in the process of subfunctionalization (Andersson et al. 2015). Revealing the responses of the remaining six related McarORs (McarORs 8, 10, 13, 22, 39, and 56; see Figure 1) is needed to conclusively inform the evolutionary history of these OR functions; in particular in this context, it could be enlightening to search for GLV-responding ORs among the orphan McarORs. The clade containing the curculionid 2-PE receptors is also intriguing due to the presence of orthologues in numerous additional species from several beetle families, including *Anoplophora glabripennis* (Cerambycidae), *R. ferrugineus*(Curculionidae), *Leptinotarsa decemlineata* (Chrysomelidae), *Tribolium castaneum* (Tenebrionidae), *Nicrophorus vespilloides* (Silphidae), and *Onthophagus taurus* (Scarabaeidae) (Antony et al. 2021; Mitchell et al. 2020). Unraveling the functions of the orthologues in these species should inform how widely conserved the ORs for 2-PE are across the Coleoptera. In moths, it was shown that OR orthologs detecting the flower compound phenylacetaldehyde were functionally conserved across eleven species from several families of the Glossata suborder although the majority of the other investigated orthologues were functionally divergent, even among related species (M. Guo et al. 2021).

The strong functional conservation among the examined OR orthologues suggests that 2-PE and GLV alcohols convey important fitness-related information to the three curculionids. Indeed, GLVs are abundant in the leaves of angiosperms and less so in conifers, and are regarded as non-host cues that typically inhibit the attraction of conifer-feeding bark beetles to their aggregation pheromones (reviewed in Zhang & Schlyter 2004). Hence, GLVs are likely used by the beetles to avoid colonizing angiosperm trees in which they cannot reproduce (Dickens et al. 1992; Schiebe et al. 2011). A role of GLVs in host choice has also been proposed for *H. abietis* (Kännaste et al. 2013; Pettersson et al. 2008).

Also 2-PE is tightly connected to the chemical ecologies - and potentially reproductive fitness - of the three investigated species. In *H. abietis* it is a potent anti-feedant, present in the gut of the beetle and in non-host plants (Axelsson et al. 2017; Eriksson et al. 2008). Through deposition of feces containing 2-PE over the laid eggs, the compound may contribute to reducing egg predation by conspecifics (Axelsson et al. 2017; Borg-Karlson et al. 2006). It is also present in the guts of both *D. ponderosae* and *I. typographus*. 2-PE inhibits attraction to the aggregation pheromone in the former species, suggesting it may contribute to termination of aggregation and induction of dispersal (Pureswaran et al. 2000). In contrast, 2-PE appears to have no effect on pheromone attraction in *I. typographus* (Schlyter et al. 1987); however, it is part of the odor blends released by several species of ophiostomatoid symbiotic fungi, which are attractive to beetles in laboratory bioassays (Kandasamy et al. 2019). These fungi, inoculated by beetles inside their galleries under the tree bark, are likely to benefit beetles by providing nutrients and through metabolism of the tree’s chemical defenses (Kandasamy et al. 2019; Kandasamy et al. 2021).

Since the ORs underlie the responses of the OSNs in the insect antennae, it is of interest to compare OR responses from *in vitro* heterologous systems with those of putatively corresponding OSN classes. Among the study species, *I. typographus* provides the best example due to the extensive electrophysiological studies that have been conducted (Andersson et al. 2009; Kandasamy et al. 2019; Kandasamy et al. 2021; Schiebe et al. 2019; Tømmerås 1985). In relation to the GLV alcohols, one OSN class (named ‘GLV-OH’) that responds most strongly and with similar sensitivity to *E*2-hexenol, *Z*3-hexenol, and 1-hexanol has been identified in *I. typographus* (Andersson et al. 2009; Kandasamy et al. 2019). This response profile resembles the ones from the GLV-responsive ORs characterized here, although several additional compounds elicited weaker secondary responses in the OSN. These secondary compounds include six-carbon aldehydes, eight-carbon alcohols as well as 2-PE (a detailed OR/OSN comparison is shown in Supplementary Table 4). The fact that 2-PE is one of the secondary compounds for the GLV-responsive OSN class may support the above-mentioned scenario that higher specificity in the extant ORs towards GLVs or 2-PE may have evolved from a more broadly tuned ancestral receptor. The three OSN-active GLVs have similar inhibitory effects on pheromone attraction of *I. typographus* and the use of a single compound (e.g., 1-hexanol) can replace a three-component GLV mixture at an equivalent release rate without compromising the inhibitory effect (Unelius et al. 2014; Zhang & Schlyter 2003). This effect on the behavior may be explained by the rather indiscriminate response of ItypOR5 to several structurally similar GLV compounds (Andersson et al. 2009; Raffa et al. 2016). Indeed, *I. typographus* has no other known OSN class that primarily responds to GLVs, although these compounds also activate OSNs primarily tuned to the less volatile compounds 3-octanol and 1-octen-3-ol (OSN class ‘C8an’) (Andersson et al. 2009; Andersson et al. 2012b). In *D. ponderosae*, coupled gas chromatographic-electroantennographic detection (GC-EAD) demonstrated antennal detection of the GLV alcohols that activate DponOR9 (Huber et al. 2000; Wilson et al. 1996). The strongest inhibitory effect on pheromone attraction was observed for *Z*3-hexenol and *E*2-hexenol (Wilson et al. 1996), that is, the two compounds that elicited the strongest responses in DponOR9. However, since (to our knowledge) no SSRs have been performed, it remains unknown whether *D. ponderosae* also has only one OSN class that responds most strongly to six-carbon GLV compounds. Likewise, SSR studies testing GLV compounds in *H. abietis* appear to be missing in the published literature (Wibe et al. 1997). In contrast to *I. typographus*, beetles feeding on angiosperms typically possess several different OSN classes primarily tuned to GLVs, each with their unique response specificity (Andersson et al. 2012a; Carrasco et al. 2019; Hansson et al. 1999; Larsson et al. 2001).

*I. typographus* also has an OSN class that primarily responds to 2-PE and secondarily to 2-phenethyl acetate and a few more compounds with weaker activity, including 1-hexanol (Kandasamy et al. 2019). Similar to ItypOR5, the secondary OSN responses were not evident in any of the three 2-PE receptors characterized in the present study (Supplementary Table 4). Higher response specificities in ORs when tested in HEK293 cells as compared to those seen in putatively corresponding OSN classes have been observed also previously, such as for ItypOR46 and ItypOR49, responding to the beetle-produced compounds ipsenol and ipsdienol, respectively (Yuvaraj et al. 2021). The reasons for the discrepancies in specificity remain unknown but could potentially be due to lower sensitivity of the HEK cell assay as compared to SSR or a consequence of the “unnatural” cellular environment in HEK cells, which may affect protein folding and hence access of ligands to their binding sites (see also Hou et al. 2020; Yuvaraj et al. 2022).

In conclusion, we report the functional characterization of six ORs from three species of the Curculionidae family, including the first characterized ORs from the devastating forest pests *H. abietis* and *D. ponderosae*. We reveal highly conserved responses to ecologically relevant odors within the two groups of assayed OR orthologues, suggesting that the detection of 2-PE and GLVs is important for the fitness of conifer-feeding curculionids. The characterized ORs were shown to be evolutionarily related with the 2-PE receptor in *M. caryae*, sharing a common ancestral OR protein. Our findings demonstrating conserved responses among beetle ORs from two taxonomic superfamilies provide new insight into the functional evolution of the OR family in this large insect order.

## Acknowledgements

We thank Göran Nordlander for providing pine weevils for RNA extractions. We thank Rikard Unelius, Suresh Ganji, Erika Wallin, Blanka Kalinová, Fredrik Schlyter, Anna Jirošová, and Wittko Francke for providing compounds. Caroline Isaksson is acknowledged for hosting of DP. The study was funded by grants from the Swedish Research Councils FORMAS (grant numbers 217-2014-689 and 2018-01444 to MNA; grant number 2018-01630 to JKY) and VR (grant number 2017-03804 to CL), the Crafoord Foundation (to MNA), the Carl Trygger Foundation (grant number CTS 17:25 to MNA), the Royal Physiographic Society in Lund (to RER, TB, and MNA), the Foundation in Memory of Oscar and Lili Lamm (to MNA), and the Max Planck Society (to EG-W and BSH).

## Data Accessibility and Benefit-Sharing

Data accessibility: The *H. abietis* RNAseq reads have been deposited in the SRA database at NCBI under the BioProject accession number PRJNA783427 (Powell, 2021). Sequences of cloned OR genes ItypOR5, ItypOR6, HabiOR3, and HabiOR4 have been deposited in GenBank under the accession numbers OL865310-OL865313 (Roberts et al., 2021b). Raw data from all HEK cell assays are reported in the supplementary materials of this article. Benefit-Sharing: Not applicable

## Author contributions

MNA conceived the project. RER and MNA conceptualized and designed the study. RER, TB, JKY, and MNA performed molecular work. RER, TB, and JKY performed cell culturing and experimental assays. EG-W, BSH, and MNA coordinated the sequencing of *H. abietis.*EG-W and DP performed transcriptome assemblies, with DP assessing OR gene expression and completeness of final assemblies. MNA annotated the HabiORs, constructed the OR phylogeny, and performed statistical analysis of the HEK cell data. MNA, JKY, and CL contributed to project supervision. MNA and RER drafted the manuscript. All authors contributed to the final version of the manuscript, and have read and approved it for submission.

## References

Andersson, M. N., Corcoran, J. A., Zhang, D.-D., Hillbur, Y., Newcomb, R. D., & Löfstedt, C. (2016). A sex pheromone receptor in the Hessian fly *Mayetiola destructor* (Diptera, Cecidomyiidae). Frontiers in Cellular Neuroscience, 10, 212.

Andersson, M. N., Grosse-Wilde, E., Keeling, C. I., Bengtsson, J. M., Yuen, M. M., Li, M., Hillbur, Y., Bohlmann, J., Hansson, B. S., & Schlyter, F. (2013). Antennal transcriptome analysis of the chemosensory gene families in the tree killing bark beetles, *Ips typographus* and *Dendroctonus ponderosae* (Coleoptera: Curculionidae: Scolytinae). BMC Genomics, 14(1), 198.

Andersson, M. N., Keeling, C. I., & Mitchell, R. F. (2019). Genomic content of chemosensory genes correlates with host range in wood-boring beetles (*Dendroctonus ponderosae*, *Agrilus planipennis*, and *Anoplophora glabripennis*). BMC Genomics, 20(1), 690.

Andersson, M. N., Larsson, M. C., & Schlyter, F. (2009). Specificity and redundancy in the olfactory system of the bark beetle *Ips typographus:* Single-cell responses to ecologically relevant odors. Journal of Insect Physiology, 55(6), 556–567.

Andersson, M. N., Larsson, M. C., Svensson, G. P., Birgersson, G., Rundlöf, M., Lundin, O., Lankinen, Å., & Anderbrant, O. (2012a). Characterization of olfactory sensory neurons in the white clover seed weevil, *Apion fulvipes* (Coleoptera: Apionidae). Journal of Insect Physiology, 58, 1325–1333.

Andersson, M. N., Löfstedt, C., & Newcomb, R. D. (2015). Insect olfaction and the evolution of receptor tuning. Frontiers in Ecology and Evolution, 3, 53.

Andersson, M. N., Schlyter, F., Hill, S. R., & Dekker, T. (2012b). What reaches the antenna? How to calibrate odor flux and ligand-receptor affinities. Chemical Senses, 37, 403–420.

Antony, B., Johny, J., Montagné, N., Jacquin-Joly, E., Capoduro, R., Cali, K., Persaud, K., Al-Saleh, M. A., & Pain, A. (2021). Pheromone receptor of the globally invasive quarantine pest of the palm tree, the red palm weevil (*Rhynchophorus ferrugineus*). Molecular Ecology, 30, 2025–2039.

Antony, B., Soffan, A., Jakše, J., Abdelazim, M. M., Aldosari, S. A., Aldawood, A. S., & Pain, A. (2016). Identification of the genes involved in odorant reception and detection in the palm weevil *Rhynchophorus ferrugineus*, an important quarantine pest, by antennal transcriptome analysis. BMC Genomics, 17(1), 69.

Axelsson, K., Konstanzer, V., Rajarao, G. K., Terenius, O., Seriot, L., Nordenhem, H., Nordlander, G., & Borg-Karlson, A.-K. (2017). Antifeedants produced by bacteria associated with the gut of the pine weevil *Hylobius abietis*. Microbial Ecology, 74(1), 177–184.

Benton, R. (2015). Multigene family evolution: perspectives from insect chemoreceptors. Trends in Ecology & Evolution, 30(10), 590–600.

Bin, S.-Y., Qu, M.-Q., Pu, X.-H., Wu, Z.-Z., & Lin, J.-T. (2017). Antennal transcriptome and expression analyses of olfactory genes in the sweetpotato weevil *Cylas formicarius*. Scientific Reports, 7(1), 11073.

Birgersson, G., Schlyter, F., Löfqvist, J., & Bergström, G. (1984). Quantitative variation of pheromone components in the spruce bark beetle *Ips typographus* from different attack phases. Journal of Chemical Ecology, 10(7), 1029–1055.

Bohbot, J. D., Jones, P. L., Wang, G., Pitts, R. J., Pask, G. M., & Zwiebel, L. J. (2011). Conservation of indole responsive odorant receptors in mosquitoes reveals an ancient olfactory trait. Chemical Senses, 36(2), 149–160.

Bolger, A. M., Lohse, M., & Usadel, B. (2014). Trimmomatic: a flexible trimmer for Illumina sequence data. Bioinformatics, 30(15), 2114–2120.

Borg-Karlson, A.-K., Nordlander, G., Mudalige, A., Nordenhem, H., & Unelius, C. R. (2006). Antifeedants in the feces of the pine weevil *Hylobius abietis*: identification and biological activity. Journal of Chemical Ecology, 32(5), 943–957.

Brand, P., & Ramírez, S. R. (2017). The evolutionary dynamics of the odorant receptor gene family in corbiculate bees. Genome Biology and Evolution, 9(8), 2023–2036.

Brand, P., Robertson, H. M., Lin, W., Pothula, R., Klingeman, W. E., Jurat-Fuentes, J. L., & Johnson, B. R. (2018). The origin of the odorant receptor gene family in insects. eLife, 7, e38340.

Butterwick, J. A., del Mármol, J., Kim, K. H., Kahlson, M. A., Rogow, J. A., Walz, T., & Ruta, V. (2018). Cryo-EM structure of the insect olfactory receptor Orco. Nature, 560(7719), 447–452.

Capella-Gutiérrez, S., Silla-Martínez, J. M., & Gabaldón, T. (2009). trimAl: a tool for automated alignment trimming in large-scale phylogenetic analyses. Bioinformatics, 25(15), 1972–1973.

Carrasco, D., Nyabuga, F. N., Anderbrant, O., Svensson, G. P., Birgersson, G., Lankinen, Å., Larsson, M. C., & Andersson, M. N. (2019). Characterization of olfactory sensory neurons in the red clover seed weevil, *Protapion trifolii* (Coleoptera: Brentidae) and comparison to the closely related species *P. fulvipes*. Journal of Insect Physiology, 119, 103948.

Clyne, P. J., Warr, C. G., Freeman, M. R., Lessing, D., Kim, J., & Carlson, J. R. (1999). A novel family of divergent seven-transmembrane proteins: candidate odorant receptors in *Drosophila*. Neuron, 22, 327–338.

Corcoran, J. A., Jordan, M. D., Carraher, C., & Newcomb, R. D. (2014). A novel method to study insect olfactory receptor function using HEK293 cells. Insect Biochemistry and Molecular Biology, 54, 22–32.

Corcoran, J. A., Sonntag, Y., Andersson, M. N., Johanson, U., & Löfstedt, C. (2018). Endogenous insensitivity to the Orco agonist VUAA1 reveals novel olfactory receptor complex properties in the specialist fly *Mayetiola destructor*. Scientific Reports, 8(1), 3489.

Dahanukar, A., Hallem, E. A., & Carlson, J. R. (2005). Insect chemoreception. Current Opinion in Neurobiology, 15(4), 423–430.

Dekel, A., Pitts, R. J., Yakir, E., & Bohbot, J. D. (2016). Evolutionarily conserved odorant receptor function questions ecological context of octenol role in mosquitoes. Scientific Reports, 6(1), 37330.

Dickens, J. C., Billings, R. F., & Payne, T. L. (1992). Green leaf volatiles interrupt aggregation pheromone response in bark beetles infesting southern pines. Experientia, 48(5), 523–524.

Eriksson, C., Månsson, P. E., Sjödin, K., & Schlyter, F. (2008). Antifeedants and feeding stimulants in bark extracts of ten woody non-host species of the pine weevil, *Hylobius abietis*. Journal of Chemical Ecology, 34(10), 1290–1297.

Gonzalez, F., Bengtsson, J. M., Walker, W. B., Sousa, M. F., Cattaneo, A. M., Montagné, N., de Fouchier, A., Anfora, G., Jacquin-Joly, E., & Witzgall, P. (2015). A conserved odorant receptor detects the same 1-indanone analogs in a tortricid and a noctuid moth. Frontiers in Ecology and Evolution, 3, 131.

Grabherr, M. G., Haas, B. J., Yassour, M., Levin, J. Z., Thompson, D. A., Amit, I., Adiconis, X., Fan, L., Raychowdhury, R., & Zeng, Q. (2011). Full-length transcriptome assembly from RNA-Seq data without a reference genome. Nature Biotechnology, 29(7), 644.

Guo, H., Huang, L.-Q., Gong, X.-L., & Wang, C.-Z. (2021). Comparison of functions of pheromone receptor repertoires in *Helicoverpa armigera* and *Helicoverpa assulta* using a Drosophila expression system. Insect Biochemistry and Molecular Biology, 141, 103702.

Guo, M., Du, L., Chen, Q., Feng, Y., Zhang, J., Zhang, X., Tian, K., Cao, S., Huang, T., Jacquin-Joly, E., et al. (2021). Odorant receptors for detecting flowering plant cues are functionally conserved across moths and butterflies. Molecular Biology and Evolution, 38, 1413–1427.

Haas, B. J., Papanicolaou, A., Yassour, M., Grabherr, M., Blood, P. D., Bowden, J., Couger, M. B., Eccles, D., Li, B., Lieber, M., et al. (2013). *De novo* transcript sequence reconstruction from RNA-seq using the Trinity platform for reference generation and analysis. Nature Protocols, 8(8), 1494–1512.

Hansson, B. S., Larsson, M. C., & Leal, W. S. (1999). Green leaf volatile-detecting olfactory receptor neurones display very high sensitivity and specificity in a scarab beetle. Physiological Entomology, 24(2), 121–126.

Hansson, B. S., & Stensmyr, M. C. (2011). Evolution of insect olfaction. Neuron, 72(5), 698–711.

Hou, X., Zhang, D.-D., Yuvaraj, J. K., Corcoran, J. A., Andersson, M. N., & Löfstedt, C. (2020). Functional characterization of odorant receptors from the moth *Eriocrania semipurpurella:* a comparison of results in the *Xenopus* oocyte and HEK cell systems. Insect Biochemistry and Molecular Biology, 117, 103289.

Hou, X.-Q., Yuvaraj, J. K., Roberts, R. E., Zhang, D.-D., Unelius, C. R., Löfstedt, C., & Andersson, M. N. (2021). Functional evolution of a bark beetle odorant receptor clade detecting monoterpenoids of different ecological origins. Molecular Biology and Evolution, 38(11), 4934–4947.

Huber, D. P. W., Gries, R., Borden, J. H., & Pierce Jr, H. D. (2000). A survey of antennal responses by five species of coniferophagous bark beetles (Coleoptera: Scolytidae) to bark volatiles of six species of angiosperm trees. Chemoecology, 10(3), 103–113.

Ji, T., Xu, Z., Jia, Q., Wang, G., & Hou, Y. (2021). Non-palm plant volatile α-pinene is detected by antenna-biased expressed odorant receptor 6 in the *Rhynchophorus ferrugineus* (Olivier)(Coleoptera: Curculionidae). Frontiers in Physiology, 1187.

Jones, P. L., Pask, G. M., Rinker, D. C., & Zwiebel, L. J. (2011). Functional agonism of insect odorant receptor ion channels. Proceedings of the National Academy of Sciences USA, 108(21), 8821–8825.

Kandasamy, D., Gershenzon, J., Andersson, M. N., & Hammerbacher, A. (2019). Volatile organic compounds influence the interaction of the Eurasian spruce bark beetle *(Ips typographus)* with its fungal symbionts. The ISME Journal, 13, 1788–1800.

Kandasamy, D., Zaman, R., Nakamura, Y., Zhao, T., Hartmann, H., Andersson, M. N., Hammerbacher, A., & Gershenzon, J. (2021). Bark beetles locate fungal symbionts by detecting volatile fungal metabolites of host tree resin monoterpenes. bioRxiv, doi:10.1101/2021.1107.1103.450988.

Katoh, K., Misawa, K., Kuma, K. i., & Miyata, T. (2002). MAFFT: a novel method for rapid multiple sequence alignment based on fast Fourier transform. Nucleic Acids Research, 30(14), 3059–3066.

Katoh, K., & Standley, D. M. (2013). MAFFT multiple sequence alignment software version 7: improvements in performance and usability. Molecular Biology and Evolution, 30(4), 772–780.

Kännaste, A., Zhao, T., Lindström, A., Stattin, E., Långström, B., & Borg-Karlson, A.-K. (2013). Odors of Norway spruce (Picea abies L.) seedlings: differences due to age and chemotype. Trees, 27(1), 149–159.

Larsson, M. C., Domingos, A. I., Jones, W. D., Chiappe, M. E., Amrein, H., & Vosshall, L. B. (2004). Or83b encodes a broadly expressed odorant receptor essential for *Drosophila* olfaction. Neuron, 43(5), 703–714.

Larsson, M. C., Leal, W. S., & Hansson, B. S. (2001). Olfactory receptor neurons detecting plant odours and male volatiles in *Anomala cuprea* beetles (Coleoptera: Scarabaeidae). Journal of Insect Physiology, 47(9), 1065–1076.

Leufvén, A., Bergström, G., & Falsen, E. (1984). Interconversion of verbenols and verbenone by identified yeasts isolated from the spruce bark beetle *Ips typographus*. Journal of Chemical Ecology, 10(9), 1349–1361.

Li, W., & Godzik, A. (2006). Cd-hit: a fast program for clustering and comparing large sets of protein or nucleotide sequences. Bioinformatics, 22(13), 1658–1659.

Manni, M., Berkeley, M. R., Seppey, M., & Zdobnov, E. M. (2021). BUSCO: Assessing genomic data quality and beyond. Current Protocols, 1(12), e323.

McKenna, D. D., Scully, E. D., Pauchet, Y., Hoover, K., Kirsch, R., Geib, S. M., Mitchell, R. F., Waterhouse, R. M., Ahn, S.-J., & Arsala, D. (2016). Genome of the Asian longhorned beetle (*Anoplophora glabripennis*), a globally significant invasive species, reveals key functional and evolutionary innovations at the beetle-plant interface. Genome Biology, 17(1), 227.

Mitchell, R. F., & Andersson, M. N. (2020). Olfactory genomics of the Coleoptera. In G. J. Blomquist & R. G. Vogt (Eds.), Insect Pheromone Biochemistry and Molecular Biology 2nd Edition (pp. 547–590). Oxford: Academic Press.

Mitchell, R. F., Hughes, D. T., Luetje, C. W., Millar, J. G., Soriano-Agatón, F., Hanks, L. M., & Robertson, H. M. (2012). Sequencing and characterizing odorant receptors of the cerambycid beetle *Megacyllene caryae*. Insect Biochemistry and Molecular Biology, 42, 499–505.

Mitchell, R. F., Schneider, T. M., Schwartz, A. M., Andersson, M. N., & McKenna, D. D. (2020). The diversity and evolution of odorant receptors in beetles (Coleoptera). Insect Molecular Biology, 29, 77–91.

Nei, M., Niimura, Y., & Nozawa, M. (2008). The evolution of animal chemosensory receptor gene repertoires: roles of chance and necessity. Nature Reviews Genetics, 9(12), 951–963.

Pettersson, M., Kännaste, A., Lindström, A., Hellqvist, C., Stattin, E., Långström, B., & Borg-Karlson, A.-K. (2008). Mini-seedlings of *Picea abies* are less attacked by *Hylobius abietis* than conventional ones: is plant chemistry the explanation? Scandinavian Journal of Forest Research, 23(4), 299–306.

Powell, D. (2021). *Hylobius abietis* antennal transcriptome. Sequence Read Archive (SRA); BioProject accession PRJNA783427.

Price, M. N., Dehal, P. S., & Arkin, A. P. (2010). FastTree 2-approximately maximum-likelihood trees for large alignments. PLoS ONE, 5(3), e9490.

Pureswaran, D. S., Gries, R., Borden, J. H., & Pierce Jr, H. D. (2000). Dynamics of pheromone production and communication in the mountain pine beetle, *Dendroctonus ponderosae* Hopkins, and the pine engraver, *Ips pini* (Say)(Coleoptera: Scolytidae). Chemoecology, 10(4), 153–168.

Raffa, K. F., Andersson, M. N., & Schlyter, F. (2016). Chapter one-Host selection by bark beetles: Playing the odds in a high-stakes game. In C. Tittiger & G. J. Blomquist (Eds.), Advances in Insect Physiology (Vol. 50, pp. 1–74). Oxford: Academic press.

Rambaut, A. (2014). FigTree v1.4.0, a graphical viewer of phylogenetic trees. http://tree.bio.ed.ac.uk/software/figtree/

Roberts, R. E., Yuvaraj, J. K., & Andersson, M. N. (2021a). Codon optimization of insect odorant receptor genes may increase their stable expression for functional characterization in HEK293 cells. Frontiers in Cellular Neuroscience, 15, 744401.

Roberts, R. E., Biswas, T., Yuvaraj, J. K., Grosse-Wilde, E., Powell, D., Hansson, B. S., Löfstedt, C., & Andersson, M. N. (2021b). Odorant receptor orthologues in weevils (Coleoptera, Curculionidae) display conserved responses to ecologically relevant odors. GenBank; accessions OL865310-OL865313.

Ruel, D. M., Yakir, E., & Bohbot, J. D. (2019). Supersensitive odorant receptor underscores pleiotropic roles of indoles in mosquito ecology. Frontiers in Cellular Neuroscience, 12, 533.

Sato, K., Pellegrino, M., Nakagawa, T., Vosshall, L. B., & Touhara, K. (2008). Insect olfactory receptors are heteromeric ligand-gated ion channels. Nature, 452(7190), 1002–1006.

Schiebe, C., Blaženec, M., Jakuš, R., Unelius, C. R., & Schlyter, F. (2011). Semiochemical diversity diverts bark beetle attacks from Norway spruce edges. Journal of Applied Entomology, 135(10), 726–737.

Schiebe, C., Unelius, C. R., Ganji, S., Binyameen, M., Birgersson, G., & Schlyter, F. (2019). Styrene, (+)-*trans*-(1*R*,4*S*,5*S*)-4-thujanol and oxygenated monoterpenes related to host stress elicit strong electrophysiological responses in the bark beetle *Ips typographus*. Journal of Chemical Ecology, 45, 474–489.

Schlyter, F., Birgersson, G., Byers, J. A., Löfqvist, J., & Bergström, G. (1987). Field response of spruce bark beetle, *Ips typographus*, to aggregation pheromone candidates. Journal of Chemical Ecology, 13(4), 701–716.

Schoville, S. D., Chen, Y. H., Andersson, M. N., Benoit, J. B., Bhandari, A., Bowsher, J. H., Brevik, K., Cappelle, K., Chen, M.-J. M., Childers, A. K., et al. (2018). A model species for agricultural pest genomics: the genome of the Colorado potato beetle, *Leptinotarsa decemlineata* (Coleoptera: Chrysomelidae). Scientific Reports, 8(1), 1931.

Shin, S., Clarke, D. J., Lemmon, A. R., Moriarty Lemmon, E., Aitken, A. L., Haddad, S., Farrell, B. D., Marvaldi, A. E., Oberprieler, R. G., & McKenna, D. D. (2018). Phylogenomic data yield new and robust insights into the phylogeny and evolution of weevils. Molecular Biology and Evolution, 35(4), 823–836.

Stensmyr, M. C., Dweck, H. K. M., Farhan, A., Ibba, I., Strutz, A., Mukunda, L., Linz, J., Grabe, V., Steck, K., Lavista-Llanos, S., et al. (2012). A conserved dedicated olfactory circuit for detecting harmful microbes in *Drosophila*. Cell, 151(6), 1345–1357.

Sullivan, B. T. (2005). Electrophysiological and behavioral responses of *Dendroctonus frontalis* (Coleoptera: Curculionidae) to volatiles isolated from conspecifics. Journal of Economic Entomology, 98(6), 2067–2078.

Tømmerås, B. Å. (1985). Specialization of the olfactory receptor cells in the bark beetle *Ips typographus* and its predator *Thanasimus formicarius* to bark beetle pheromones and host tree volatiles. Journal of Comparative Physiology A, 157(3), 335–342.

Unelius, R. C., Schiebe, C., Bohman, B., Andersson, M. N., & Schlyter, F. (2014). Non-host volatile blend optimization for forest protection against the European spruce bark beetle, *Ips typographus*. PLoS ONE, 9(1), e85381.

Vosshall, L., Amrein, H., Morozov, P., Rzhetsky, A., & Axel, R. (1999). A spatial map of olfactory receptor expression in the *Drosophila* antenna. Cell, 96, 725–736.

Wang, X., Wang, S., Yi, J., Li, Y., Liu, J., Wang, J., & Xi, J. (2020). Three host plant volatiles, hexanal, lauric acid, and tetradecane, are detected by an antenna-biased expressed odorant receptor 27 in the dark black chafer *Holotrichia parallela*. Journal of Agricultural and Food Chemistry, 68(28), 7316–7323.

Wibe, A., Borg-Karlson, A.-K., Norin, T., & Mustaparta, H. (1997). Identification of plant volatiles activating single receptor neurons in the pine weevil (*Hylobius abietis*). Journal of Comparative Physiology A, 180(6), 585–595.

Wicher, D., Schäfer, R., Bauernfeind, R., Stensmyr, M. C., Heller, R., Heinemann, S. H., & Hansson, B. S. (2008). *Drosophila* odorant receptors are both ligand-gated and cyclic-nucleotide-activated cation channels. Nature, 452(7190), 1007–1011.

Wilson, I. M., Borden, J. H., Gries, R., & Gries, G. (1996). Green leaf volatiles as antiaggregants for the mountain pine beetle, *Dendroctonus ponderosae* Hopkins (Coleoptera: Scolytidae). Journal of Chemical Ecology, 22(10), 1861–1875.

Xie, J., Liu, T., Yi, C., Liu, X., Tang, H., Sun, Y., Shi, W., Khashaveh, A., & Zhang, Y. (2022). Antenna-biased odorant receptor HvarOR25 in *Hippodamia variegata* tuned to allelochemicals from hosts and habitat involved in perceiving preys. Journal of Agricultural and Food Chemistry, 70, 1090–1100.

Yuvaraj, J. K., Andersson, M. N., Corcoran, J. A., Anderbrant, O., & Löfstedt, C. (2018). Functional characterization of odorant receptors from *Lampronia capitella* suggests a non-ditrysian origin of the lepidopteran pheromone receptor clade. Insect Biochemistry and Molecular Biology, 100, 39–47.

Yuvaraj, J. K., Jordan, M. D., Zhang, D.-D., Andersson, M. N., Löfstedt, C., Newcomb, R. D., & Corcoran, J. A. (2022). Sex pheromone receptors of the light brown apple moth, *Epiphyas postvittana*, support a second major pheromone receptor clade within the Lepidoptera. Insect Biochemistry and Molecular Biology, 141, 103708.

Yuvaraj, J. K., Roberts, R. E., Sonntag, Y., Hou, X., Grosse-Wilde, E., Machara, A., Zhang, D.-D., Hansson, B. S., Johanson, U., Löfstedt, C., et al. (2021). Putative ligand binding sites of two functionally characterized bark beetle odorant receptors. BMC Biology, 19, 16.

Zhang, Q.-H., & Schlyter, F. (2003). Redundancy, synergism, and active inhibitory range of non-host volatiles in reducing pheromone attraction in European spruce bark beetle *Ips typographus*. Oikos, 101(2), 299–310.

Zhang, Q.-H., & Schlyter, F. (2004). Olfactory recognition and behavioural avoidance of angiosperm nonhost volatiles by conifer-inhabiting bark beetles. Agricultural and Forest Entomology, 6(1), 1–19.

