## Supplemental Figure 1 for "Odorant receptor orthologues in conifer-feeding beetles display conserved responses to ecologically relevant odors"

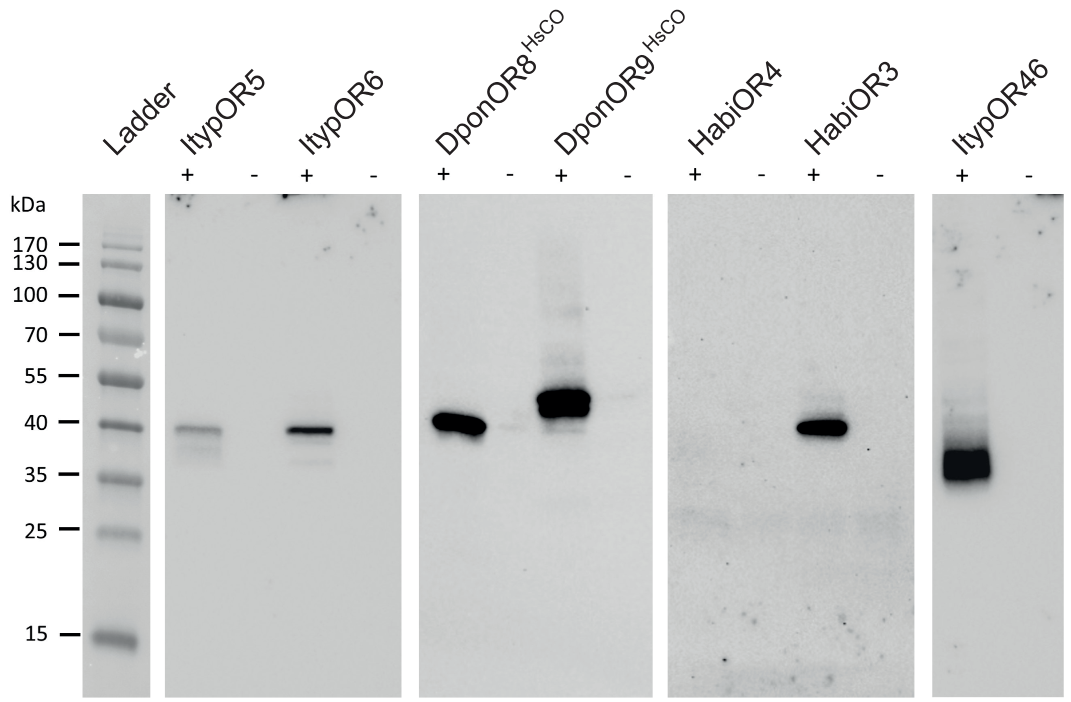


**Supplementary Figure 1.** Results from Western blot analyses of V5-tagged odorant receptors (ORs) from TREx/HEK293 cells. Included are ORs from *Ips typographus* (ItypOR5 and ItypOR6), *Dendroctonus ponderosae* (DponOR8 and DponOR9, expressed from *Homo sapiens* codon-optimized, 'HsCO', genes), and *Hylobius abietis* (HabiOR3 and HabiOR4). Proteins were only detected from cells induced (+) to express the OR genes, and not from non-induced (−) control cells, indicating proper regulation by the repression system. ItypOR46 was included as a positive control (see Yuvaraj et al., 2021).
