## Supplementary material for "Odorant receptor orthologues in conifer-feeding beetles display conserved responses to ecologically relevant odors": Supplemantal Table 2

**Supplementary Table 2.** Compounds used for characterization of odorant receptors (ORs), with their purities and source information. The green leaf volatiles that activate HabiOR4/DponOR9/ItypOR5 alongside the most structurally similar inactive compounds are highlighted in green. 2-Phenylethanol that activates HabiOR3/DponOR8/ItypOR6 and structurally similar inactive compounds are highlighted in purple.

| **Compound** | **Purity (%)** | **Source**** |
| --- | --- | --- |
| Acetophenone | 99 | Acros |
| 4-Allylanisole | >99 | Aldrich |
| Amitinol | 91 | R. U. |
| Benzaldehyde | >99 | Kebo |
| Benzyl acetate | >99 | Aldrich |
| Benzyl alcohol | 99 | Aldrich |
| (±)-*exo*-Brevicomin | 99 | W. F. |
| (±)-*endo*-Brevicomin | 96 | Synergy Semiochemicals |
| (±)-Camphene | 95 | Aldrich |
| (±)-Camphor | 97 | Aldrich |
| (+)-3-Carene | 99 | Aldrich |
| (±)-Carvone | >99 | Fluka |
| 1,8-Cineole | 99 | Aldrich |
| (5*S*,7*S*)-*trans*-Conophthorin | 94 | W. F. |
| *p*-Cymene | >99 | Acros |
| 3,4-Dimethoxytoluene | 98 | Givaudan-Roure |
| 4-Ethylguaiacol | 98 | Sigma-Aldrich |
| Eugenol methyl ether | >99 | Fluka |
| (±)-Frontalin | >99 | Synergy Semiochemicals |
| Geranyl acetate | 97 | Aldrich |
| Geranylacetone | >99 | Fluka |
| (±)-Grandisol (grandlure I) | 95 | Bedoukian (E. W.) |
| Hexanal* | 96 | Sigma |
| 1-Hexanol | >99 | Fluka |
| *E*2-Hexenal* | 98 | Aldrich |
| *E*2-Hexenol | 96 | Aldrich |
| *E*3-Hexenol* | 98 | Aldrich |
| *Z*2-Hexenol* | 95 | Aldrich |
| *Z*3-Hexenol | 98 | Aldrich |
| Indole | >99 | Aldrich |
| (±)-Ipsdienol | 94 | Bedoukian |
| (±)-Ipsenol | 95 | Synergy Semiochemicals |
| α-Isophorone | >99 | Acros |
| (+)-Isopinocamphone | >99 | R. U. |
| (−)-Isopinocamphone | >99 | R. U. |
| Lanierone | >99 | Synergy Semiochemicals |
| 4-Methylanisole | >99 | Fluka |
| 2-Methyl-3-buten-2-ol | >99 | Acros |
| (±)-2-Methylbutyl acetate | >99 | SAFC |
| 3-Methylbutyl acetate | 97 | Sigma-Aldrich |
| Myrcene | 95 | Sigma-Aldrich |
| *E*-Myrcenol | >99 | Fytofarm |
| (±)-Myrtenol | 96 | G. B. |
| Nonanal | 98 | Acros |
| (±)-3-Octanol | 97 | Sigma-Aldrich |
| (±)-1-Octen-3-ol | 98 | Janssen Chimica |
| 2-Phenethyl acetate | >99 | Aldrich |
| 2-Phenylethanol | >99 | Sigma |
| (+)-α-Pinene | 98 | Janssen Chimica |
| (−)-α-Pinene | >99 | Fluka |
| (+)-Pinocamphone | 84 | R. U |
| (−)-Pinocamphone | 81 | R. U. |
| (±)-Sabinene | 97 | Chemos GmbH |
| Styrene | >99 | Fluka |
| γ-Terpinene | 97 | Aldrich |
| Terpinolene | 98 | Fluka |
| (+)-*trans*-4-Thujanol | 97 | Sigma-Aldrich |
| (4*S*)-*cis*-Verbenol | 99 | Borregaard |
| (+)-*trans*-Verbenol | 92 | SCM |
| (−)-*trans*-Verbenol | 97 | SciTech Ltd., Prague |
| (−)-Verbenone | >99 | Fluka |
| 4-Vinylanisole | 97 | Aldrich |
| VUAA1 | 98 | Sigma-Aldrich |

*Compound tested only on green leaf volatile-responding ORs (ItypOR5, DponOR9, HabiOR4).

**Abbrevations: R. U. = gift from Rikard Unelius (Linnaeus University, Kalmar, Sweden); E. W. = gift from Erika Wallin (Mid Sweden University, Sweden); W. F. = gift from Wittko Francke (University of Hamburg, Germany); G. B. = gift from Gunnar Bergström (University of Gothenburg, Sweden).
