## Supplemental Table 4 for "Odorant receptor orthologues in conifer-feeding beetles display conserved responses to ecologically relevant odors"

**Supplementary Table 4.** Comparing responses of *Ips typographus* odorant receptors (ItypORs) in HEK cells with responses from putatively corresponding olfactory sensory neuron (OSN) classes previously shown to respond primarily to green leaf volatile alcohols (GLV-OHs) or 2-phenylethanol (2-PE), respectively.

|  | **Receptor** | **Neuron class^1,2^** | **Receptor** | **Neuron class^2^** |
| --- | --- | --- | --- | --- |
| **Stimulus** | ItypOR5 | GLV-OH | ItypOR6 | 2-PE |
| 1-Hexanol | 2 | 1 |  | 4 |
| *Z*3-Hexenol | 2 | 1 |  |  |
| *E*2-Hexenol | 1 | 1 |  |  |
| Hexanal |  | 2 |  |  |
| *E*2-Hexenal |  | 3 |  |  |
| (±)-1-Octen-3-ol |  | 3 |  |  |
| (±)-3-Octanol |  | 4 |  |  |
| 2-Phenylethanol |  | 3 | 1 | 1 |
| 2-Phenethyl acetate |  |  |  | 2 |
| Benzyl alcohol |  | 3 |  | 3 |
| (±)-*exo*-Brevicomin |  | 4 |  |  |

- Numbers in the table reflect the rank order between compounds in terms of response magnitude in the respective *in vitro* or *in vivo* system (1 = strongest response, 2 = second strongest etc.; no number = compound inactive). Compounds eliciting similar responses have the same number. Only compounds tested on both the OR and OSNs are listed.
- OSN data from **^1^**Andersson et al. (2009) and **^2^**Kandasamy et al. (2019).
